# Focused Ultrasound Augments the Delivery and Penetration of Model Therapeutics into Cerebral Cavernous Malformations

**DOI:** 10.1101/2024.08.27.609060

**Authors:** Delaney G. Fisher, Matthew R. Hoch, Catherine M. Gorick, Claire Huchthausen, Victoria R. Breza, Khadijeh A. Sharifi, Petr Tvrdik, G. Wilson Miller, Richard J. Price

## Abstract

**BACKGROUND:** Cerebral cavernous malformations (CCMs) are vascular neoplasms in the brain that can cause debilitating symptoms. Current treatments pose significant risks to some patients, motivating the development of new nonsurgical options. We recently discovered that focused ultrasound-microbubble treatment (FUS) arrests CCM formation and growth. Here, we build on this discovery and assess the ability of FUS to deliver model therapeutics into CCMs.

**METHODS:** Quantitative T1 mapping MRI sequences were used with 1 kDa (MultiHance; MH) and 17 kDa (GadoSpin D; GDS) contrast agents to assess the FUS-mediated delivery and penetration of model small molecule drugs and biologics, respectively, into CCMs of Krit1 mutant mice.

**RESULTS:** FUS elevated the rate of MH delivery to both the lesion core (4.6-fold) and perilesional space (6.7-fold). Total MH delivery more than doubled in the lesion core and tripled in the perilesional space when FUS was applied immediately prior to MH injection. For the model biologic drug (i.e. GDS), FUS was of greater relative benefit, resulting in 21.7-fold and 3.8-fold delivery increases to the intralesional and perilesional spaces, respectively

**CONCLUSIONS:** FUS augments the delivery and penetration of therapeutics into the complex and disorganized CCM microenvironment. Benefits to small molecule drug delivery are more evident in the perilesional space, while benefits to biologic delivery are more evident in CCM cores. These findings, when combined with ability of FUS alone to control CCMs, highlight the potential of FUS to serve as a powerful non-invasive therapeutic platform for CCM.

## Introduction

Cerebral cavernous malformation (CCM) is a vascular disorder characterized by the development of abnormal, dilated clusters of blood vessels in the brain^1^. These malformations are prone to repetitive hemorrhages, inducing debilitating symptoms, such as neurological deficits, seizures, and stroke, in affected individuals^2–4^. Presently, the prevailing recourse for treating symptomatic CCMs is surgical resection. However, surgical excision of CCMs poses an elevated risk of complications and morbidity^5,6^.

Despite multiple studies investigating therapeutic targets and screening pharmacological treatments for CCM^7–18^, no approved drug treatments exist for CCM. The majority of tested pharmacological agents for CCM are small molecules. In comparison, larger biologic molecules, such as antibodies and gene therapies, have not been as thoroughly explored. Additionally, drugs showing promise in acute CCM models often demonstrate limited efficacy in more clinically-representative chronic models, suggesting a potential need for greater local doses of these therapies^19,20^. Indeed, though CCMs are known to be more permeable than healthy cerebrovasculature^21–24^, delivery of systemically administered drugs to these complex lesions is poorly understood.

Focused ultrasound-mediated blood-brain barrier opening (FUS) has emerged as a promising non-invasive drug delivery technology^25–27^. With FUS, acoustic energy is concentrated into a confined volume, facilitating the oscillation of intravenously administered gas-filled microbubbles within blood vessels of the targeted region. These microbubble oscillations induce a transient disruption of endothelial tight junctions^28^ and increased active transport^29^, enabling therapeutic delivery across the blood-brain barrier (BBB). Magnetic resonance imaging (MRI) guidance permits spatial targeting of FUS to specific brain regions and BBB opening confirmation through the accumulation of gadolinium-based MRI contrast agents.

Recently, our group demonstrated that FUS, in the absence of therapeutic delivery, arrests the formation and growth of CCMs^30^. This remarkable observation prompts the exploration of the combined impact of FUS-mediated lesion stabilization and therapeutic delivery on CCMs. While our previous study also confirmed that FUS enhanced MRI contrast agent delivery beyond the natural permeability of CCMs, the MRI sequences only provided qualitative assessments. In particular, this qualitative MRI approach was sub-optimal for visualizing contrast agent delivery to the lesion core. Indeed, the cellular and molecular composition within the lesion core, including mutated endothelium, red blood cells, and their byproducts, differs substantially from the perilesional space, characterized by dense populations of astrocytes and microglia^30,31^. This difference not only affects MRI signal but may also have important implications for drug delivery to these distinct regions. Consequently, to facilitate comprehensive measurements of potential enhanced therapeutic delivery with FUS in the intricate CCM microenvironment, quantitative MRI methods are needed.

Building on our recent observations^30^, the objective of this study was to establish a foundation for therapeutic delivery approaches that harness and synergize with this potent bioeffect. We have previously demonstrated that T1-contrast mapping can enable longitudinal, quantitative concentration measurements of gadolinium-based molecules in CCMs^31^. Thus, this is an ideal method to measure FUS-induced changes for therapeutic delivery to CCMs. To this end, we employed T1-contrast mapping MRI to quantitatively evaluate the delivery of 1 kDa and 17 kDa molecules to CCMs, comparing outcomes with and without FUS. This study lays the groundwork for treatment regimens of FUS-delivered molecules capable of inducing CCM regression and clearance.

## Results

### FUS Enhances Delivery Rate of MultiHance in CCMs

We first tested if FUS would increase the delivery rate of a model small molecule drug to the CCM microenvironment. To this end, we employed T1 mapping MRI to measure the concentration of the MRI contrast agent MultiHance (MH; gadobenate dimeglumine; ∼1 nm; ∼1 kDa) before and after the application of FUS in CCM mice. One frontal hemisphere received FUS (n=6 sonication targets) with passive cavitation detection (PCD) feedback control 20 minutes after intravenous (i.v.) MH injection. During FUS, peak-negative pressures (PNPs) settled into a range of 0.3 to 0.4 MPa (**Figure 1A**), yielding integrated acoustic emissions shown in **Figure 1B**. The contralateral hemisphere was not sonicated (i.e., FUS^-^ control) to illustrate baseline CCM permeability. As expected, prior to FUS, CCMs in the non-sonicated and sonicated hemispheres displayed similar rates of MH accumulation (**Figure 1C, D, F**). After FUS, the rate of MH accumulation in the lesion core was enhanced (**Figure 1D**), increasing to well-above (4.6-fold) the rate of MH accumulation in FUS^-^ CCMs (p=0.0221; **Figure 1E**). Predictably, the perilesional space of these CCMs also displayed the same permeability rate prior to FUS in both groups (**Figure 1F**). FUS then increased perilesional MH delivery rate by 6.7-fold over the rate of MH accumulation in FUS^-^ CCMs (p<0.0001; **Figure 1G**). These results indicate that FUS enhances the delivery rate of a model small molecule drug to both the lesion core and the surrounding CCM microenvironment.

**Figure 1.**
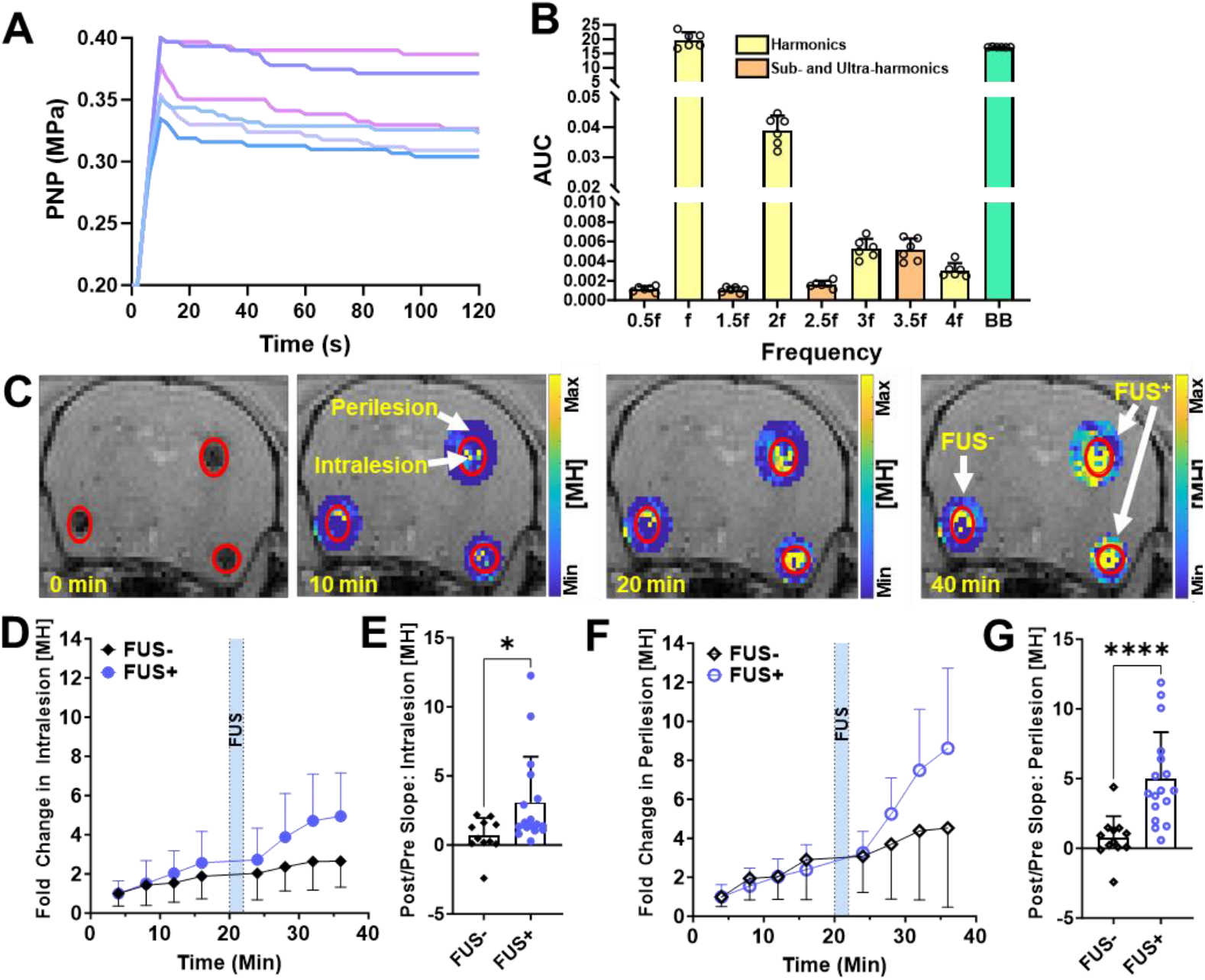
FUS Enhances Delivery Rate of MultiHance to CCMs. **A, B)** PNP histories (**A**) and integrated acoustic emissions (**B**) for FUS treatments (n=6). f = fundamental frequency; BB = broadband. **C)** T1 mapping MRIs illustrating MH accumulation, in and around 3 CCMs, over a 40 min time period. Perilesional and intralesional regions are denoted. FUS was applied to 2 of the 3 CCMs at the 20 min timepoint. A marked increase in MH concentration is evident in and around FUS-treated CCMs at 40 min. **D**) Temporal fold change in intralesional MH concentration over the average initial concentration for FUS^-^ and FUS^+^ CCMs. FUS^+^ CCMS were treated at 20 min after MH injection (blue shading). **E**) Slope ratios (Post-FUS/Pre-FUS) derived from intralesional MH concentration plots. *p=0.0221; Mann-Whitney test. **F**) Temporal fold change in perilesional MH concentration over the average initial concentration for FUS^-^ and FUS^+^ CCMs. **G**) Slope ratios (Post-FUS/Pre-FUS) derived from perilesional MH concentration plots. ****p<0.0001; Mann-Whitney test.

### FUS Enhances Total Delivery of MultiHance in CCMs

We then tested the ability of FUS to augment model small molecule drug delivery to CCMs when applied concurrently with model drug injection, reflecting the clinical staging to maximize overall delivery. On day 1, T1 mapping MRI was conducted on CCM mice following i.v. MH injection to measure baseline permeability (**Figure 2A**). On day 2, FUS was applied to one frontal hemisphere of the same CCM mice immediately before i.v. MH injection. T1 mapping MRI was conducted for 20 mins thereafter (**Figure 2B**). FUS markedly boosted the intralesional MH delivery rate, as well as mean intralesional MH concentration (p=0.0070; **Figure 2C**), with a 2.5-fold enhancement evident at 20 minutes. Area under the curve (AUC) analysis, representing the integrated exposure of CCM tissue to the model drug through time, indicates that FUS enhances intralesional model drug exposure by 1.9-fold (p=0.0122; **Figure 2D**). Regarding the perilesional space, MH concentration was also markedly elevated with FUS (p=0.0005; **Figure 2E**), with a 3.1-fold enhancement evident at 20 minutes. AUC yielded a 2.9-fold increase in model drug exposure over FUS^-^ CCMs (p=0.0007; **Figure 2F**). Notably, MH delivery after FUS becomes evident in the perilesional space (**Figure 2E**) before the intralesional space (**Figure 2C**) (0.040 mM versus 0.029 mM, respectively, after 5 minutes), yet both locations plateau to the same mean concentration by 20 mins post-injection (0.069 mM each). These results reveal that FUS can more than double the amount of a small molecule delivered to the lesion core and triple the amount in the surrounding CCM microenvironment.

**Figure 2.**
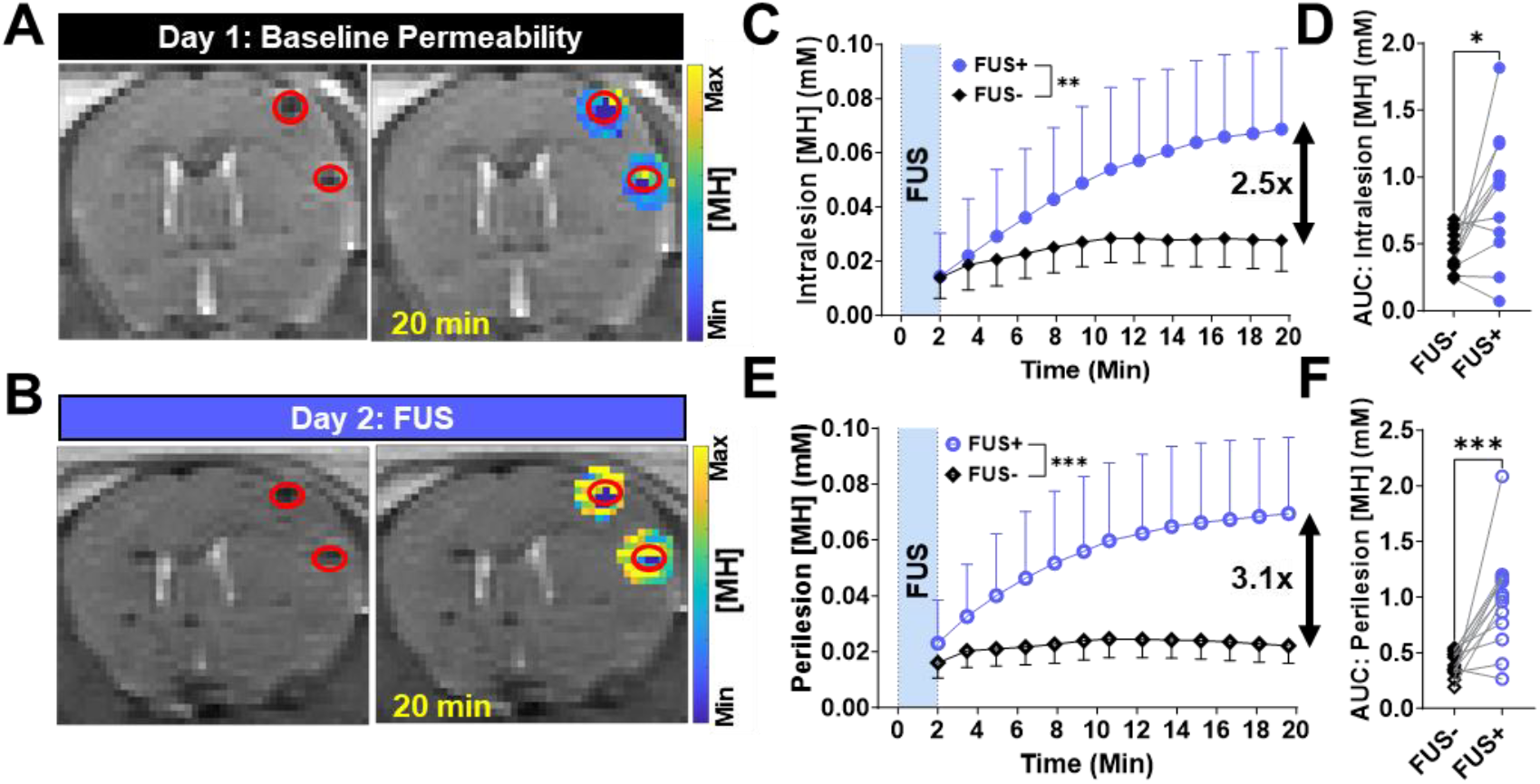
FUS Enhances Total Delivery of MultiHance to CCMs. **A)** T1 mapping MRI illustrating baseline permeability to MH, in and around 2 CCMs, 20 min after MH injection on day 1. **B)** T1 mapping MRI illustrating enhanced MH accumulation, in and around 2 CCMs, 20 min after MH injection and FUS treatment on day 2. **C**) Intralesion MH concentration as a function of time after MH injection. Baseline permeability to MH (FUS^-^) was measured on day 1, with FUS (FUS^+^) measurements made on paired CCMs on day 2. **p=0.0070; Repeated measures two-way ANOVA with Geisser-Greenhouse correction. **D**) Area under the curve (AUC) metric derived from intralesion concentration data in C. *p=0.0122; Wilcoxon matched-pairs signed rank test. **E**) Perilesion MH concentration as a function of time after MH injection. ***p=0.0005; Repeated measures two-way ANOVA with Geisser-Greenhouse correction. **F**) Area under the curve (AUC) metric derived from perilesional concentration data in E. ***p=0.0007; Wilcoxon matched-pairs signed rank test.

### FUS Enhances Total Delivery of GadoSpin D in CCMs

Next, we tested the potential for FUS to enhance the total delivery and penetration of a biologic, which are typically >1 kDa, to CCMs. To this end, we employed the MRI contrast agent GadoSpin D (GDS; dendritic Gd-chelate; ∼5 nm; ∼17 kDa) as a model biologic. As in the MH experiments (**Figure 2**), baseline permeability of CCMs to GDS was measured on day 1 (**Figure 3A**). On day 2, FUS was applied to paired CCMs from day 1. FUS improved total GDS delivery in both the intralesional and perilesional spaces compared to baseline CCM permeability (**Figure 3B**). FUS elicited a striking increase in GDS delivery to the lesion core (p=0.0106; **Figure 3C**), reaching 21.7-fold at 20 minutes. AUC was increased 4.8-fold in CCM cores with FUS (p=0.0078; **Figure 3D**). Meanwhile, perilesional delivery of GDS was also enhanced with FUS (p= 0.0021; **Figure 3E**), reaching a 3.8-fold increase at 20 minutes. For GDS in the perilesional space, integrated tissue-drug exposure increased 2.2-fold (p=0.0195; **Figure 3F**). The lesion core and perilesional space followed a similar temporal pattern of GDS enhancement following FUS, but the intralesional space peaked at a higher concentration than the perilesional space (0.010 mM versus 0.0076 mM, respectively).

**Figure 3.**
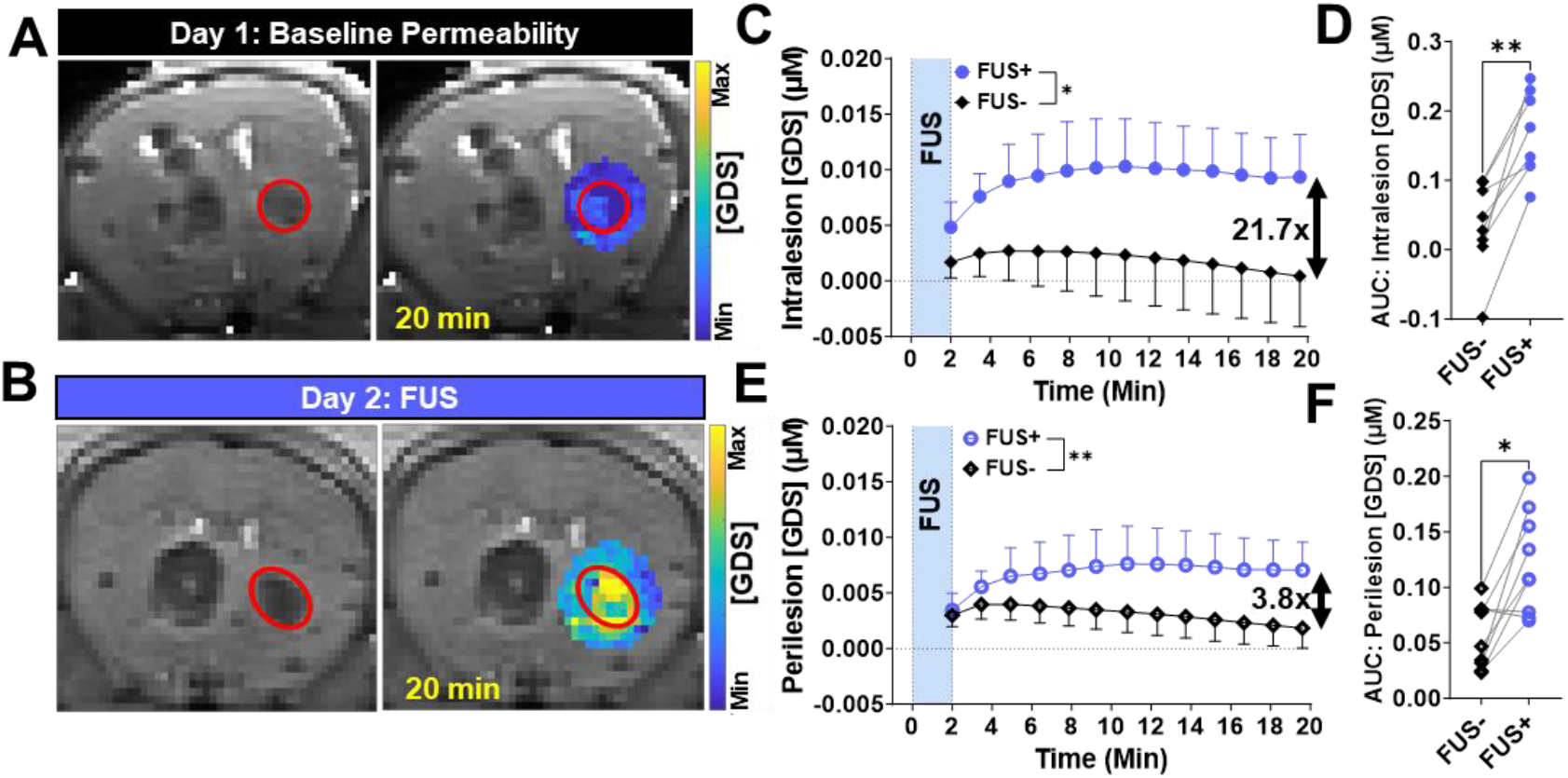
FUS Enhances Total Delivery of Gadospin D in CCMs. **A)** T1 mapping MRI illustrating baseline permeability to GDS, in and around a CCM, 20 min after GDS injection on day 1. **B)** T1 mapping MRI illustrating enhanced GDS accumulation, in and around a CCM, 20 min after GDS injection and FUS treatment on day 2. **C**) Intralesion GDS concentration as a function of time after GDS injection. Baseline permeability to GDS (FUS^-^) was measured on day 1, with FUS (FUS^+^) measurements made on paired CCMs on day 2. *p=0.0106; Repeated measures two-way ANOVA with Geisser-Greenhouse correction. **D**) Area under the curve (AUC) metric derived from intralesion concentration data in C. **p=0.0078; Wilcoxon matched-pairs signed rank test. **E**) Perilesion GDS concentration as a function of time after GD injection. **p=0.0021; Repeated measures two-way ANOVA with Geisser-Greenhouse correction. **F**) Area under the curve (AUC) metric derived from perilesional concentration data in E. *p=0.0195; Wilcoxon matched-pairs signed rank test.

### Comparison of FUS-Mediated MultiHance and GadoSpin D Delivery to Intralesion and Perilesion CCM Compartments

We also investigated whether FUS differentially affects the delivery of MH and GDS to intralesional and perilesional regions of CCMs. To this end, we first needed to verify that the applied FUS PNP, as well as the resultant MB activity, were equivalent in the MH and GDS experiments. The PNP histories for the MH (**Figure 4A**) and GDS (**Figure 4B**) experiments followed similar trajectories, and there were no differences in average (**Figure 4C**) and maximum (**Figure 4D**) applied PNP. Moreover, MB activity, as assessed by acoustic emissions across several key spectral domains (i.e. sub-harmonic, harmonic, ultra-harmonic, and broadband), was equivalent for the MH and GDS experiments. Thus, any differences between MH and GDS delivery were not due to differences in FUS application and/or MB response.

**Figure 4.**
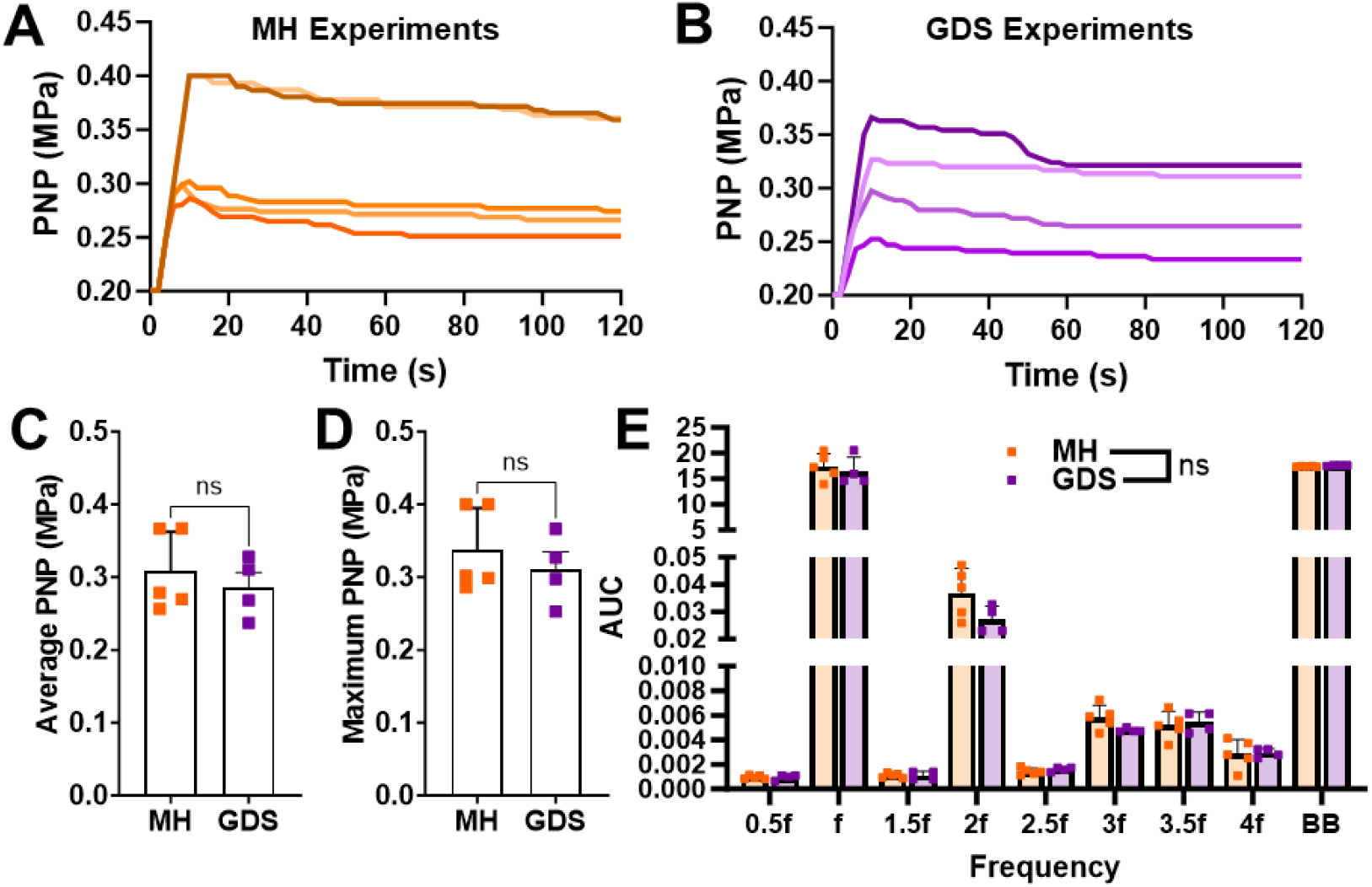
Focused Ultrasound Application in Multihance and Gadospin D Delivery Experiments was Comparable. **A, B)** PNP histories during BBB opening by acoustic emissions feedback control for MH (n=5) (A) and GDS (n=4) (B) treatments. **C, D**) Average (C) and maximum (D) PNPs for MH and GDS delivery experiments. Mann-Whitney tests. **E**) Integrated acoustic emissions from key spectral domains for MH and GDS delivery experiments. f = fundamental frequency; BB = broadband. Mann-Whitney tests.

When comparing GDS to MH delivery using the AUC metric, similar levels of FUS-mediated delivery enhancement (i.e. FUS^+^/FUS^-^) to both the intralesional (**Figure 5A**) and perilesional spaces (**Figure 5B**) were observed, with GDS exhibiting a slight trend (p=0.23) over MH in intralesional AUC augmentation (**Figure 5A**). To then examine whether intralesional or perilesional AUC augmentation might be favored for one or both of the contrast agents, we calculated the ratio of intralesional FUS-mediated AUC enhancement over perilesional FUS-mediated AUC enhancement. Resultant values >1 suggest greater relative intralesional amplification (**Figure 5C**). By this metric, GDS exhibited greater relative FUS-mediated augmentation of delivery to the intralesional space when compared to MH (**Figure 5C**). We then repeated this analysis using maximum concentration as the key metric. As with the AUC comparisons, there was no difference between the 2 contrast agents with respect to FUS-mediated intralesional (**Figure 5D**) and perilesional (**Figure 5E**) delivery augmentation, but there was greater relative amplification of delivery to the intralesional space for GDS (**Figure 5F**). Finally, we compared post-FUS times to maximum concentration in the intralesional and perilesional spaces for MH and GDS (**Figure 5G and 5H**). For both regions, GDS reached its maximum concentration in about 10 min after FUS, while MH concentration was typically still increasing at the final (20 min) timepoint.

**Figure 5.**
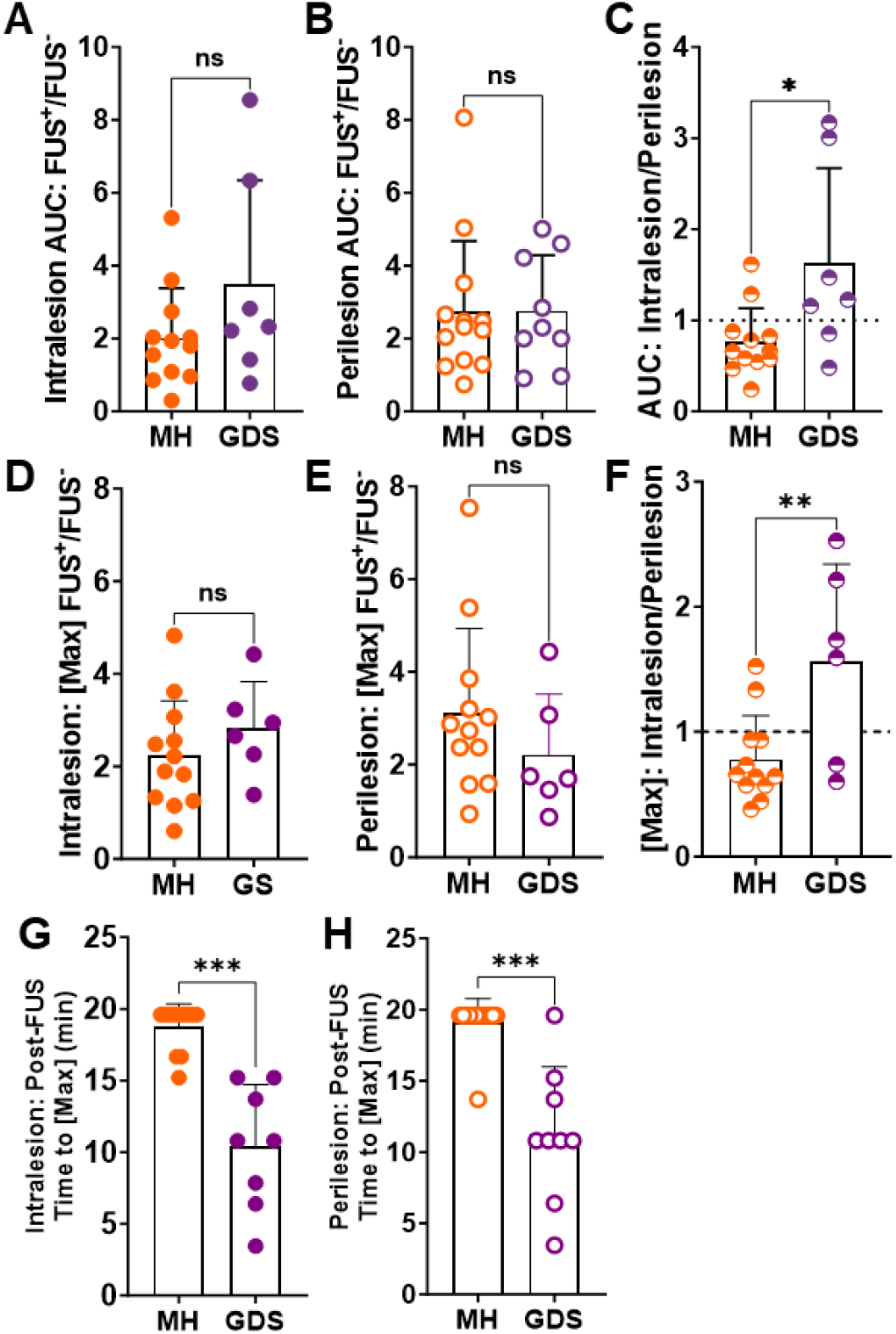
Comparison of FUS-Mediated MultiHance and GadoSpin D Delivery to Intralesion and Perilesion CCM Compartments. **A, B**) Intralesion (A) and perilesion (B) AUC ratios (FUS^+^/FUS^-^) for MH and GDS. Mann-Whitney tests. **C)** Intralesion/perilesion ratios of AUC ratios for MH and GDS. *p=0.045; Mann-Whitney test. **D, E)** Intralesion (D) and peilesion (E) maximum concentration ratios (FUS^+^/FUS^-^) for MH and GDS. Mann-Whitney tests. **F)** Intralesion/perilesion ratios of maximum concentration ratios for MH and GDS. *p=0.025; Mann-Whitney test. **G, H**) Intralesion (G) and perilesion (H) post-FUS time to maximum concentration for MH and GDS. ***p<0.001; Mann-Whitney tests.

## Discussion

We previously elucidated that FUS can arrest CCM growth and formation, even in the absence of therapeutic delivery^30^. Here, we aimed to advance the synergistic potential for concurrent therapeutic delivery with this approach. Utilizing longitudinal T1 mapping MRI, we quantified the impact of FUS on therapeutic delivery of model small molecule drugs and biologics to CCMs. Our findings revealed a significant enhancement in the delivery rate of a 1 kDa small molecule, exhibiting a 4.6-fold increase in the lesion core and a 6.7-fold increase in the perilesional space. Moreover, FUS augmented overall delivery of both the 1 kDa small molecule and a 17 kDa model biologic to CCMs, with a 2.5-fold increase for the model small molecule drug and an impressive 22-fold increase for the model biologic in the lesion core. In the perilesional space, there was a 3.1-fold increase for the model small molecule drug and a 3.8-fold increase for the model biologic. GDS reached its post-FUS maximum concentration sooner than MH, suggesting there may be a more transient delivery window for biologics. Finally, our analysis uncovered a nuanced aspect of FUS enhancement, wherein the relative FUS-mediated effect is more pronounced for the small molecule in the perilesional space and for the model biologic in the lesion core. These results collectively establish a robust foundation for employing FUS in targeted therapeutic delivery regimens to effectively mitigate CCMs.

### T1 Mapping MRI Enables Spatiotemporal, Intra-CCM, Delivery Comparisons

Given the notable heterogeneity in baseline CCM permeability^23,31,32^, methods allowing for comparative measurements in the same CCMs over time are important for generating statistical power and robust conclusions. We have previously shown that T1 mapping MRI enables longitudinal and quantitative assessments of contrast agent deposition in individual CCMs^31^. Thus, it was reasonable to leverage this MRI approach to measure model drug delivery to CCMs with FUS. Yet another advantage of T1 mapping MRI is that it has sufficient spatial resolution to discern differences in discrete CCM tissue compartments. Indeed, the lesion core harbors mutated, cavernous vessels filled with clotted blood components, while the perilesional space surrounds the core with dense populations of astrocytes, microglia, and macrophages^30,31,33^. These regional differences in the CCM microenvironment pose varying biotransport challenges that can influence the efficacy of different delivery approaches and molecule sizes. T1 mapping MRI enabled us to measure the exact concentration of MH and GDS in both the intralesional and perilesional spaces of the CCM microenvironment, both with and without FUS.

### Differential Spatial Delivery Augmentation for Varying-Sized Molecules with FUS

One unexpected and potentially important finding that arose from our spatiotemporally detailed T1 mapping results was that FUS differentially augments the delivery of small and large molecules to the two pre-defined CCM tissue compartments (i.e. lesion core vs. perilesional space). Specifically, FUS provided a greater relative benefit for (i) model small molecule drug delivery to the perilesional space and (ii) model biologic delivery to the lesion core. This effect is evident when using either AUC (**Figure 5C**) or maximum concentration (**Figure 5F**) as the metric of interest. To explore the potential causes behind the observed differential spatial delivery of varying-sized molecules with FUS, we first emphasize that FUS is known to offer varying degrees of benefit based on the transport properties of a given molecule^34^. Noting that the increase in permeability induced by FUS had a greater effect for MH in the perilesional space, we postulate that the benefit of FUS for small molecule drug delivery in regions with an already disrupted BBB (e.g. the lesion core) is less than in areas that have a more intact BBB (e.g. the perilesional space). Conversely, for a larger molecule like GDS (17 kDa; 5 nm), crossing the disrupted BBB in the lesion core may be less feasible due to biophysical constraints limiting the transport of a larger molecule. FUS partially alleviates these constraints, ultimately providing more relative benefit for larger molecules than for small molecules in the already leaky CCM core. In perilesional regions harboring a more intact BBB, even small molecules cannot effectively cross into the brain parenchyma. Thus, FUS yields a larger benefit for small molecule delivery in this region. Moreover, for larger molecules, the advantage of FUS may be less pronounced in regions with a previously intact BBB than in regions with a previously disrupted BBB, once again due to increased biophysical transport constraints.

We also note that differences in BBB closure time, as well as clearance mechanisms within the CCM microenvironment, for small molecules and biologics could impact the integrated exposure of tissue to drug. Here, GDS reached its maximum concentration at ∼10 minutes after FUS (**Figure 5G and 5H**), while MH concentration was often still increasing at 20 min after FUS. This is consistent with the hypothesis that the BBB in and around CCMs closes fairly rapidly to larger therapeutics, which could factor into how injections are timed with respect to FUS application. Regarding clearance, while there is evidence that FUS alters clearance mechanisms through modification of the glymphatic system^35–37^ and BBB efflux pumps^38,39^, its specific influence on the clearance of varying-sized molecules remains unclear. Our data indicate that GDS concentrations rapidly decrease without FUS when compared to MH without FUS or GDS with FUS, highlighting that differential clearance is also likely a significant determinant of tissue-drug exposure.

### Potential for Clinical Impact on Therapeutic Delivery in CCM

Here, we demonstrate that FUS enhances therapeutic delivery for molecules of different sizes in both the CCM core and surrounding perilesional space. In the clinic, this will translate to increased local delivery for any given standard systemic dose, thereby increasing therapeutic index. Furthermore, enhanced on-target drug delivery reduces the risk of side effects associated with off-target delivery. The greater benefit observed for larger molecules with FUS opens the door for biologic delivery exploration for CCM. Indeed, our study highlights that, in the absence of FUS, the delivery of a 5 nm model biologic drug (GDS) is minimal. There also may be rapid clearance from both the intralesional and perilesional spaces. However, with FUS, biologic-sized molecules are more effectively retained in both CCM compartments. These findings pave the way for future investigation into even larger agents with promising therapeutic potential for CCM, such as antibodies and gene therapy vectors.

Notably, FUS also offers a level of precision that can be customized for either familial or sporadic cases of CCM. In these studies, we induce BBB opening in a substantial volume—almost one-quarter—of the CCM brain. In contrast, our previous study showcased targeting FUS to a smaller volume of the CCM brain^30^. For patients, FUS can be tailored to target a large volume, which may be necessary for familial patients with multiple CCMs, or it can be focused on a singular CCM, as would be needed for sporadic cases. Moreover, the region of delivery can also be adapted for the mechanism of action of the delivered therapeutic. Drugs with a preventative effect could be more widely delivered than those with specific corrective functions in the CCM microenvironment. Ultimately, given its ability to stabilize lesions and seamlessly integrate with therapeutic delivery, FUS may offer a powerful platform for the treatment of CCM via image-guided drug and gene delivery.

## Materials and Methods

### Animals

All animal experiments adhered to ethical guidelines and were approved by the University of Virginia Animal Care and Use Committee. The animals were housed in accordance with standard laboratory conditions, maintaining a temperature of 22°C and a 12-hour light/12-hour dark cycle. The generation of the CCM murine model was established as previously detailed^31^. Briefly, *Krit1*^fl/null^ or *Krit1*^fl/fl^ male or females were generated under the endothelial promoter Pdgfb^CreER^. On postnatal day 5, induction of *Krit1* was initiated with a subcutaneous injection of tamoxifen (50 µL at 2mg/mL in corn oil). Genotypes were subsequently verified using Transnetyx (Cordova, TN). Mice were studied between 2 and 3 months old.

### MRI Acquisition

Data for T1 maps were acquired with a set of multi-slice 2D spin echo (SE) images at varied repetition times (TR) to generate a saturation recovery curve. 2 sets of 7 images, for a total of 14 scans, were acquired prior to FUS and contract agent administration to obtain saturation recovery curves with a satisfactory dynamic range. The two sets of image series were offset by the slice thickness in the slice select plane to ensure 3D coverage of the brain. The parameters for these scans were: TR=790, 1040, 1350, 1750, 2300, 3215, and 7000 ms, TE=6.71 ms, slice thickness=0.6 mm, slice gap=0.6 mm, FOV=35 × 35 mm, matrix size=180 × 180, rare factor=10, and R= 0.194 × 0.194 × 0.6 mm^3^. After FUS and contrast agent administration, 14 SE images were acquired with identical parameters except at a fixed TR=1040 ms. The acquisitions alternated between slice package orientations resulting in 7 images at each slice profile geometry. Time per acquisition was 1 minute and 28 seconds.

### Data Processing

A saturation recovery approach was utilized to calculate M_0_ and all T1 values (pre and post contrast) on a voxel-by-voxel basis by fitting the data to the signal equation:

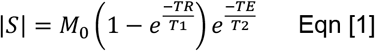

In equation 1, |*S*| is the magnitude of the signal within the voxel, *M*_0_ is the product of the thermal equilibrium magnetization and coil sensitivity, TR is the repetition time (ms), T1 is the spin-lattice relaxation (ms), TE is the echo time (ms), and T2 is the spin-spin relaxation (ms). The echo time exponential is assumed to be 1 due to TE<<T2, resulting in the final form seen in equation 2.

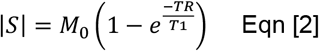

A custom MATLAB script fit the signal magnitude data on a voxel-by-voxel basis to equation 2. Each fitting procedure simultaneously fit the data to 8 functions: function 1 incorporated the 7 pre-contrast variable TR scans, while functions 2-8 incorporated the singular scan at a fixed TR but different time points. The fits were constrained to having the same *M*_0_ value but allowed different T1 values. Pre-contrast and post-contrast T1 values were then used to calculate the contrast agent concentration on a voxel-by-voxel basis at each time point using equation 3.

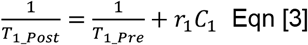

In equation 3, *T*_1 *Post*_ is the post-contrast value at a particular time point (ms), *T*_1 *Pre*_ is the pre-contrast T1 value (ms), r_1_ is the contrast agent relaxivity (L/mmol/ms), and C1 is the contrast agent concentration (mM). At the conclusion of this process, concentration values for slice package 1 existed for time points (minutes): 1.47, 4.40, 7.33, 10.27, 13.2, 16.13, and 19.07, while concentration values for slice package 2 existed for time points (minutes): 2.93, 5.87, 8.80, 11.73, 14.67, 17.60, and 20.53. To obtain 3D coverage at each time point, concentration data was calculated at the missing time points by linearly interpolating between the acquired points. This required an assumption of 0 concentration at minute 0 for slice package 2. The 20.53-minute time point was not used because it required data be extrapolated past minute 19.07 for slice package 1.

A second custom MATLAB script was used to calculate average concentrations with manually drawn regions of interest (ROIs) on the concentration maps. To ensure the iron rich intralesional data was not skewed by susceptibility artifacts, a data exclusion method was developed. Briefly, a ROI of healthy brain tissue on the contralateral hemisphere was used to calculate an average residuals value for the fit. If any residuals value for the voxels within the lesion core were 3 times greater than this average, they were excluded from the analysis. The value of 3 was empirically determined. To maintain consistency within data processing, this was also applied to all perilesional data.

### FUS Blood-Brain Barrier Opening

The FUS procedure was conducted using the RK-300 small bore FUS device (FUS Instruments, Toronto, CA). Mice were prepared by shaving and depilating their heads before being placed in a supine position and coupled to the transducer using degassed ultrasound gel. Blood-brain barrier opening was achieved using a 1.1 MHz single-element transducer with a 10 ms burst length over a 2000 ms period. A total of 60 sonications were administered during a 2-minute sonication duration. The FUS Instruments software, operating in the “Blood-brain Barrier” mode, facilitated PCD-modulated PNP. The feedback control system parameters were set as follows: a starting pressure of 0.2 MPa, pressure increment of 0.05 MPa, maximum pressure of 0.4 MPa, 20 sonication baselines without microbubbles, area under the curve (AUC) bandwidth of 500 Hz, AUC threshold of 10 standard deviations, pressure drop of 0.95, and frequency selection of the subharmonic, first ultraharmonic, and second ultraharmonic. Optison™ (GE HealthCare) microbubbles were intravenously injected as a bolus dose of 10^5 microbubbles per gram of body weight. Prior to sonication, the distribution of microbubble diameter and concentration was assessed using a Coulter counter (Multisizer 3; Beckman Coulter, Fullerton, California). T1 mapping MRI sequences were used to guided sonication targeting. Six non-overlapping sonication targets were placed over one frontal hemisphere with placement optimized to target CCMs.

### Contrast Agent Injections

MultiHance ^**®**^ (gadobenate dimeglumine; Bracco) and GadoSpin D™ (dendritic Gd-chelate; Viscover) were injected as a bolus intravenously at a dose of 0.01 and 0.0002 mmol, respectively, diluted in saline. Injection of contrast agent was given immediately prior to MRI acquisition for FUS^-^ control studies and immediately following the initiation of FUS for FUS^+^ studies.

### Passive Cavitation Detection

Acoustic emissions during FUS were detected with a fiber-optic hydrophone (Precision Acoustics, Dorset, UK) of 10 um diameter and 15 mm aperture center-mounted within the ultrasound transducer. Emissions data was processed with a custom MATLAB script. The area under the curve of the acoustic emissions at the subharmonic (0.5f) and ultra-harmonics (1.5f, 2.5f) after applying a 300 Hz bandwidth filter. Broadband emissions were evaluated by summing acoustic emissions following the removal of all emissions at the fundamental frequency, harmonics (2f, 3f, 4f), subharmonic (0.5f), and ultra-harmonics (1.5f, 2.5f, 3.5f).

### Statistical Analysis

All results reported with error bars are means with standard deviation. The “n” values per group are made evident either by individual data points shown or statement of “n” value in figure or figure legend. Statistical significance was assessed at p < 0.05 for all experiments and were calculated using GraphPad Prism 9 (San Diego, USA). Statistical tests are provided in the figure legends.

## Author Contributions

DGF, MH, and RJP conceptualized the study. DGF and MH conducted the FUS experiments with the aid of CMG in animal preparation. MRI sequences and analysis were optimized by MH and GWM. MRI data was acquired and processed by MH and analyzed by DGF and MH. KAS generated experimental animals. DGF and MH designed the figures and wrote the manuscript. GWM, PT, and RJP edited the manuscript. All authors approved the manuscript.

## Acknowledgements

This work was supported by funding from NIH R01CA279134, R01EB030409, R01EB030744, and R21NS118278 to RJP; NIH R21NS116431 and grants from Focused Ultrasound Foundation, Be Brave for Life Foundation, and Alliance to Cure Cavernous Malformation to PT; NIH R01CA226899 to GWM; and American Heart Association 830909 to DGF. MRI was performed in the University of Virginia Molecular Imaging Core Laboratory, with support for the 9.4T Bruker scanner from NIH S10OD025024. We thank Dr. Kevin Whitehead for kindly providing the mouse strains used in this study. We are also grateful to Jeremy Gatesman of the University of Virginia Center for Comparative Medicine for assistance with catheterization procedures.

## Notes

### Competing Interest Statement

The authors have declared no competing interest.

### Summary of Updates

The title has been slightly changed, with the word "impels" replaced by "augments". An error in the units for Figure 3 has been corrected. The correct concentration units are micromoles, not millimoles.

